# Generative Chemistry Platform for Small Molecules Targeting RNA: A Case Study for Chemical Optimization

**DOI:** 10.64898/2026.05.08.723908

**Authors:** Timothy E. H. Allen, Susan M. Boyd, Maurinne Bonnet, Rabia T. Khan

**Affiliations:** Serna Bio; CompChem Solutions Ltd; University of Michigan, Biointerfaces Institute

## Abstract

We introduce the Serna Bio GenAI platform, a generative chemistry and multiparametric optimization platform for the design of RNA-targeting small molecules. Targeting RNA with small molecules has proven historically challenging but offers notable potential upsides, including access to unique mechanisms of action and the ability to target otherwise untargetable genes. We consider a major challenge here to be designing chemistry specific to RNA-targeting. Molecular design is a valuable application of AI in drug discovery, but many publicly available models use training data focused on protein-targeting - the modality best historically explored in drug discovery. We showcase the difference and value in building a specifically RNA-targeting platform, comparing its performance to state-of-the-art public chemical generators and experimentally validating its chemical designs in comparison to chemistry designed by a human expert.

## Introduction

Despite significant advances in drug discovery technologies, a vast expanse of human biology remains inaccessible to classical approaches. Historically, the pharmaceutical industry has focused on the proteome [1], yet proteins represent only a small fraction of the functional genome. By contrast, RNA encodes for approximately 70% of the human genome, offering an untapped landscape for therapeutic intervention [2,3]. Targeting RNA with small molecules does more than expand the list of available targets; it enables sophisticated regulatory mechanisms that are impossible to achieve by targeting proteins alone. Through splice modulation and translational enhancement, small molecules can proactively reshape the transcriptome, allowing for the correction of genetic errors and the restoration of protein levels before a disease-state protein is ever synthesized [4].

While there is significant excitement in the field of RNA-targeting small molecules, to date, only one drug, risdiplam, with a known mechanism of action (MOA) modulating RNA function, has been approved by the FDA [5], excluding decades-old antibiotics. Several compounds are currently in clinical trials, including MYB-targeting compounds from Remix Therapeutics [6] and Rgenta Therapeutics [7], and an mHTT-targeting compound from SkyHawk Therapeutics [8], but the number of clinical RNA-targeting small molecules remains limited. Several challenges in RNA-small molecule drug discovery have been well covered previously [9], and these include challenges in optimizing RNA-targeting small molecules, challenges working in a limited-data space [10], a relatively poor understanding of the chemical matter that binds to RNA [11] and uncertainty in how binding relates to functional effects [12]. In addition, we propose that optimizing RNA-targeting small molecules is challenging due to SAR (structure-activity relationship) cliffs [13], where small changes in chemical structure remove compound activity and derail optimization efforts.

It has been established that the toolkit for targeting proteins does not work for RNA [3]. Strategies developed for targeting proteins may not be applicable to RNA, such as optimizing molecules for target binding affinity, which has been shown to be decoupled from functional activity [12]. We hypothesize that the challenges for discovery and optimization of small molecules for targeting RNA arise from the historical precedent of protein-targeting drug discovery. Designing and optimizing small molecules to target RNA may require different molecule designs and permutations, as may be required for any new modality. Machine learning architectures capable of designing novel chemistry may be of value in this task, as other machine learning models are also seeing application in RNA science [14–16].

Generative chemistry AI models enable the rapid de novo design of novel small molecules with desired properties, including molecular activity, desirable physicochemistry and synthetic tractability [17–22]. These models are powerful tools that work best in an iterative loop with human medicinal chemists to ensure the candidates generated are practical and therapeutically aligned. While a wide variety of generative chemistry models are available in the public domain, the training data for these have inevitably been influenced by protein-targeting drug discovery. For example, ChEMBL [23–27], the LINCS L1000 gene expression dataset [28–30], the GEOM-Drugs 3D chemistry set [31–35] and the ZINC 250k dataset (which contains ‘drug-like’ structures) [36–42] are all influenced by publications and patents, frequently from pharmaceutical research targeting proteins. We hypothesize that using models trained using these datasets will lead to chemical solutions tailored towards protein-targeting drug discovery processes, and it is these kinds of solutions that are finding SAR cliffs when being applied to RNA-targeting drug discovery. We believe that training or fine-tuning models on relevant RNA-small molecule data will move their focus away from proteins and into RNA-targeting chemical space and will remove human bias from design-make-test (DMT) cycles. Coupling this explicit direction towards RNA-targeting with models that effectively explore the chemical space around a seed compound, as would typically be done during drug discovery in hit-to-lead and lead optimization, could address issues with SAR cliffs, leading to improved outcomes for the designed compounds in RNA-targeting drug discovery.

To test this hypothesis, we have developed an ensemble of generative ML models trained on a proprietary dataset of more than 2.4 million RNA-small molecule interactions specifically with the aim of enabling SAR exploration. We aimed to understand if:

- (1) Generative chemistry models trained on RNA-binding chemistry design different compounds than public state-of-the-art models trained on protein-targeting datasets
- (2) Models trained on RNA-binding chemistry design more potent analogues when exploring a known example from RNA-targeting drug discovery (risdiplam)
- (3) When applied to an internal drug discovery program, we can improve compound optimization (e.g. to improve potency or solubility), by designing molecules differentiated from a classical medicinal chemistry approach

In this work, we present the Serna Bio Generative Chemistry AI platform for the design and multiparametric optimization of RNA-targeting small molecules for hit-to-lead and lead optimization. We show that our RNA-small molecule-specific generative chemistry models, trained on our corpus of Serna Bio RNA-small molecule data, designs compounds different from those designed by public chemical generators and human medicinal chemists. Furthermore, we compare the compounds designed in both *in silico* and experimental tests and show that our platform is designing highly valuable compounds for hit-to-lead and lead optimization in RNA-targeting drug discovery campaigns. Finally, we conclude by showing how our platform is driving success in an RNA-targeting drug discovery program, improving compound potency ∼3X in a single DMT cycle and designing high-potency analogues at a similar rate to a human medicinal chemist.

## Materials and Methods

### Serna Bio RNA-small molecule dataset

We have collected more than 2.4 million RNA-small molecule binding datapoints [43] to enable this work. RNA-small molecule binding data was collected using the Automated Ligand Identification System (ALIS) [44] or the self-assembled monolayer desorption ionization-affinity selection-mass spectrometry (SAMDI-ASMS) system [45]. Data was collected across 7 independent screening exercises, spanning 181,632 unique small molecules and 85 RNA targets. 5,726 (3.2%) of the small molecules bind to at least one of the RNA targets tested. Targets are specifically selected motifs within genes known to form secondary structures, including G-quadruplexes, multi-way junctions, and stem-loops.

The physicochemical properties of the compounds tested, the binders identified, and small molecules from the public R-BIND dataset [46] were calculated using RDKit [47] and are included in the supplementary data (Fig. S1-3, Table S1-3).

### Fine-tuning chemical generators

In this work, a SMILES BERT masked language model (MLM) [48] and a Matched Molecular Pairs (MMP) algorithm were fine-tuned/trained using an identical Serna Bio RNA-small molecule dataset for comparison. In selecting these models for our study, we considered and trained additional architectures but are restricting our discussion here to the SMILES BERT and MMP algorithms for brevity. Models were trained using an identical dataset of experimentally determined RNA-targeting small molecules from our 2.4 million datapoints represented as Canonicalized SMILES (SMILES canonicalized using RDKit [47]). For this task, five known RNA-binding small-molecule seed compounds were withheld from the fine-tuning dataset and only used as seeds for each model in a validation exercise. These specific molecules were excluded from model training/testing to prevent data leakage. The remaining RNA binders were provided to the models for model training/testing.

For fine-tuning SMILES BERT models, the following hyperparameters were used:

- mask_probability: 0.15
- batch_size = 8
- n_epochs = [5, 10]
- learning_rate = 0.0001
- weight_decay = 0
- test_fraction=0.2
- loss_function = CrossEntropyLoss

The model was fine-tuned using stochastic test set masking for 5 and 10 epochs using a held-out test set (20% of the training data allocated randomly). A random split of training and testing data was deemed appropriate for unsupervised model training to design new analogues within the domain of RNA-binding chemical space, allowing the model to be exposed to all scaffolds we have identified as RNA-binding in our dataset. We acknowledge that this may mean our generative models are less suitable for out-of-distribution molecule generation.

For comparison, an MMP database was constructed using RDKit using all RNA binders in the training/test set to identify valid molecular transformations between RNA-binding small molecules. For molecular generation, the following parameters were used:

- radius = 0
- min_pairs = 1

The combination of the three models trained using the Serna Bio dataset (two SMILES BERT and one MMP) and the one SMILES BERT pre-trained generative model is used as a generative ensemble in subsequent tasks as the Serna Bio GenAI platform ensemble. Molecular generation is entirely ligand-dependent, with only chemical structural information required to be fed into the generative ensemble as SMILES strings.

In the Serna Bio GenAI platform, compounds designed by the Generative Chemistry ensemble are assessed in the Predictive Model ensemble, and predictions are used as defined by a User to rank compounds for synthesis and experimental evaluation. Iterative improvement of generative and predictive models can be undertaken after each DMT cycle using an active learning loop, but for the examples explored in this work, no active learning has been deployed.

### Generating risdiplam analogues using REINVENT

Risdiplam analogues were designed using REINVENT, using the implementation of REINVENT 3.2 available through MOE [49]. The prior used was the default ChEMBL database. REINVENT-generated analogues were designed using a pharmacophore model-guided and an ECFP4 Tanimoto similarity-guided generator, each trained for 600 reinforcement steps. These were selected to give a broad spectrum of both diverse (provided by the pharmacophore model-guided generator) and similar (provided by the ECFP4 Tanimoto similarity-guided generator) compounds to the input seed compound (risdiplam), as designed by REINVENT.

### Generating risdiplam analogues using MolMIM

Risdiplam analogues were designed using MolMIM using the online portal [50] default settings for the two presented sampling algorithms (CMA-ES Controlled Generation, quantitative estimate of druglikeness (QED) Optimized, Number of Molecules to generate = 30, Similarity Constraint = 0.3, Particles = 30, Iterations = 60 and Sampling Standard Deviation, Number of Molecules to generate = 30, Sample Radius = 0.5). Note that the MolMIM CMA-ES Controlled Generation sampling algorithm is non-deterministic and will produce different samples each time it is run, even with the same inputs.

### Calculation of cheminformatics parameters

Tanimoto Similarity: All referenced Tanimoto Similarity [51] (ECFP, radius = 2, len = 2048) values were calculated using RDKit [47].

Synthetic accessibility (SA) score: SA scores [52] were calculated using sascorer in RDKit [47] to provide a simple numerical metric to estimate the ease of compound synthesis. SA scores are measured on a scale of 1 - 10, with lower scores indicating a compound is more synthesizable

### Risdiplam-SMN2-U1 duplex docking model

Solution structures of the SMN2-U1 duplex target of risdiplam in the presence (6HMO [53]) and absence (6HMI [54]) of risdiplam illustrate how the small molecule binds to the duplex structure and stabilizes a bulged adenine. For docking purposes, one of the NMR models containing reasonable interactions between the small molecule and RNA target (model 3) was identified, and risdiplam was docked into that structure based on the following constraints:

- A donor to the adjacent phosphate backbone was required
- An acceptor onto the NH2 of Adenine-14 was required
- Hydrophobic regions corresponding to the 3 hydrophobic sections of risdiplam were required

Top poses were identified similar to the NMR pose (Fig. 3b). Risdiplam analogues designed by the Serna Bio GenAI, REINVENT and MolMIM were prepared as 3D structures and passed to the GOLD docking model (GOLD 2023.3 from CCDC [55]) for virtual screening. GOLD PLP/CHEMPLP fitness scores [56] were calculated for each analogue (plotted in Fig. 3c). More rigorous physics-based binding energy calculations, including Free Energy Perturbation, Relative Binding Free Energy, and Absolute Binding Free Energy, offer potential improvements in virtual screening over the docking model outlined above, but require more precise structures of the ligand in complex with its target than were available during this study [57,58].

### High Content Imaging GLUT1 protein expression measurements

Target protein expression was quantified using high content imaging (HCI) in an 8-point concentration-response experiment. Test concentrations were 2-fold serial dilutions from a top test concentration of 10µM in DMSO (0.1% concentration in final test media). HeLa cells were treated with test compounds for 24 hours in triplicate. Antibody staining and HCI were used to quantify the level of the target protein and the count of cell nuclei as a proxy for cell viability. Image segmentation and multivariate data analysis yielded concentration response curves (Fig. 7) quantifying the percentage change in protein level normalized to 0.1% DMSO vehicle.

EC25 values were determined using CDD Vault [59] to fit a Hill Equation curve to the concentration-response data. The EC25 value is determined using an absolute 25% upregulation of the target protein compared to vehicle control.

Curve fitting (Fig. 7) was performed using Prism for an agonist vs. response with a variable slope (four parameters) using the Goodness of Fit Method, Robust Sum of Squares method.

### Quantitative Polymerase Chain Reaction GLUT1 mRNA expression measurements

Target mRNA expression was quantified using Quantitative Polymerase Chain Reaction (qPCR). HeLa cells were seeded at a density of 5000 cells/well and treated with compounds at 3 µM for 24 hours in triplicate. Lysates were prepared using the TaqMan™ Fast Advanced Cells-to-CT™ Kit (Thermo Fisher Scientific) according to the manufacturer’s instructions. Multiplex qPCR was performed using TaqMan assays for the target and B2M (housekeeping gene control). Relative gene expression was calculated via the 2-ΔΔ CT method, which is presented as fold change normalized to the DMSO control (Fig. 7). Statistical significance was determined by an unpaired t-test (ns = not significant, *p <= 0.05, **p <= 0.01, ***p <= 0.001, ****p <= 0.0001).

### Glucose uptake measurements

Glucose uptake was measured in the human microvascular endothelial cell line hCMEC/D3 cells (Sigma-Aldrich) using the Glucose uptake Glo assay (Promega). hCMEC/D3 cells were seeded at 20,000 cells per well into a 96-well plate. At 24 hours post-seeding the cells were incubated with vehicle (0.1% DMSO) or compound for 24h prior to assay readout. Glucose uptake was performed by incubating the cells for 5 min with 1mM 2-Deoxyglucose in PBS and the subsequent assay readout was performed according to the manufacturer’s instructions. Significance was determined for comparisons of each compound to the vehicle control using an unpaired t-test (ns = not significant, *p <= 0.05, **p <= 0.01, ***p <= 0.001, ****p <= 0.0001).

### ALIS biophysical binding study to GLUT1 RNA motif

RNA-small molecule binding data for the discovered GLUT1 RNA motif was conducted using the Automated Ligand Identification System (ALIS) [44] at a single test concentration. The RNA target (5μM) was incubated for 30 min at room temperature with each compound (10μM). Resulting RNA-small molecule complexes were stabilized at 4 °C and separated from unbound compounds using size exclusion chromatography (column: 2.1mm I.D. × 50mm, PolyHydroxyethyl A, 60, (PolyLC, Columbia MD); solvent: 700mM ammonium acetate, pH 8; flow rate: 0.3mL/min at 4 °C). Isolated complexes were then delivered to a reverse phase chromatography column (Targa C18, 0.5mm I.D. × 50mm, 5μm packing material (Higgins Analytical, Mountain View, CA) running at a linear gradient of 0-90% acetonitrile/0.2% formic acid in 2.5 min (total run time of 4 min including column equilibration) at 20μL/min and 60 °C) to dissociate, desalt and separate small molecules for subsequent detection using electrospray ionization-based mass spectrometry (Thermo Scientific™Exactive™ Orbitrap high-resolution, accurate-mass mass spectrometer detector operating at 100,000 resolution with a mass accuracy of <5 ppm and a scan rate of 1 Hz). Binders were selected using the following criteria: uniqueness (absence from the blank control), ASMS score > 50 (depends on mass accuracy and isotopic distribution), inspection of peak area and S/N ratio.

## Results

### The Serna Bio GenAI platform designs different compounds better suited for RNA when compared to REINVENT and MolMIM

At Serna Bio, we have fine-tuned generative chemistry models using a proprietary dataset of more than 2.4 million RNA-small molecule datapoints [43]. Our generative chemistry models are an ensemble of different ML architectures, including language and cheminformatics chemical generators. These chemical generators have been specifically selected to help us explore the chemical space around a provided seed compound - as would be required in hit-to-lead or lead optimization during drug discovery - not to ‘scaffold hop’ into different areas of chemical space. Following chemical generation, the generative chemistry platform ranks compounds using a multiparametric optimization (MPO) function, where the parameters can include a variety of desirable chemical properties (for example, RNA-binding probability, synthetic accessibility and druglikeness). As generative chemistry AI methods can design thousands of compounds, we use this MPO to rank the designs for synthesis and experimental testing.

As we move through the molecular optimization process, an active learning loop is used for continuous improvement of the platform. Different generative models and different features of the MPO are used to optimize the key characteristics needed to achieve the target product profile (TPP) for the program. We believe that using this approach can reduce hit-to-lead and lead optimization timelines from more than 40 months [60] to fewer than 20.

Our first aim was to evaluate the value of training models on our data using a known RNA-targeting small molecule seed - risdiplam [5] and by comparing the compounds designed to those designed by public generative chemistry methods. We have assessed success for compounds designed by the generators around two specific themes:

- How useful would the designed compounds be if we are trying to explore the chemical SAR around risdiplam for hit-to-lead and lead optimization in drug discovery? We consider analogues that are similar to the seed compound useful in this aspect. The generation of diverse analogues is useful in other contexts, but it is not the aim of this study.
- How well do the designed analogues dock to risdiplam’s target (the SMN2-U1 duplex [61])? Do we expect them to have higher affinity and improved potency? We would want the models to improve the potency of any provided seed compound. We use an *in silico* docking study as a proxy for this while acknowledging that docking is not the equivalent of an experimental confirmation of molecular activity.

For a model to be useful in advancing this hypothetical RNA-targeting drug discovery campaign, a generative model would need to perform well in both of these tasks, as well as generate a collection of valid, unique, and synthetically accessible small molecules.

Risdiplam analogues were generated using publicly available, state-of-the-art chemical generators REINVENT [49], designed by AstraZeneca, MolMIM [50], designed by Nvidia, and our GenAI platform. The REINVENT architecture consists of recurrent neural networks and transformer architectures for molecular generation, and MolMIM also uses transformers. REINVENT and MolMIM were run using default settings for the purpose of this comparison, and we acknowledge that tuning and objective alignment can strongly affect the outcome of generative AI models. Hence, our case study should be seen as an attempt to compare our models to these defaults, and we do not claim the results discussed here are the best that can be achieved with REINVENT and MolMIM.

Given risdiplam as a seed compound, REINVENT generated 11,481 compounds (Fig. 1). Both the Serna Bio GenAI and MolMIM generated 61 and 46 compounds, respectively. Importantly, none of the Serna Bio GenAI-designed compounds were also designed by REINVENT or MolMIM and the maximum pairwise chemical similarity between the designed compound lists, as measured using Tanimoto similarity (Fig. 2) is low for all three pairs of generators (including comparing the designs from MolMIM to REINVENT, Fig. 2c). This supports our hypothesis that our platform, and its models trained using RNA-small molecule data, is designing different chemistry than public models.

**Figure 1.**
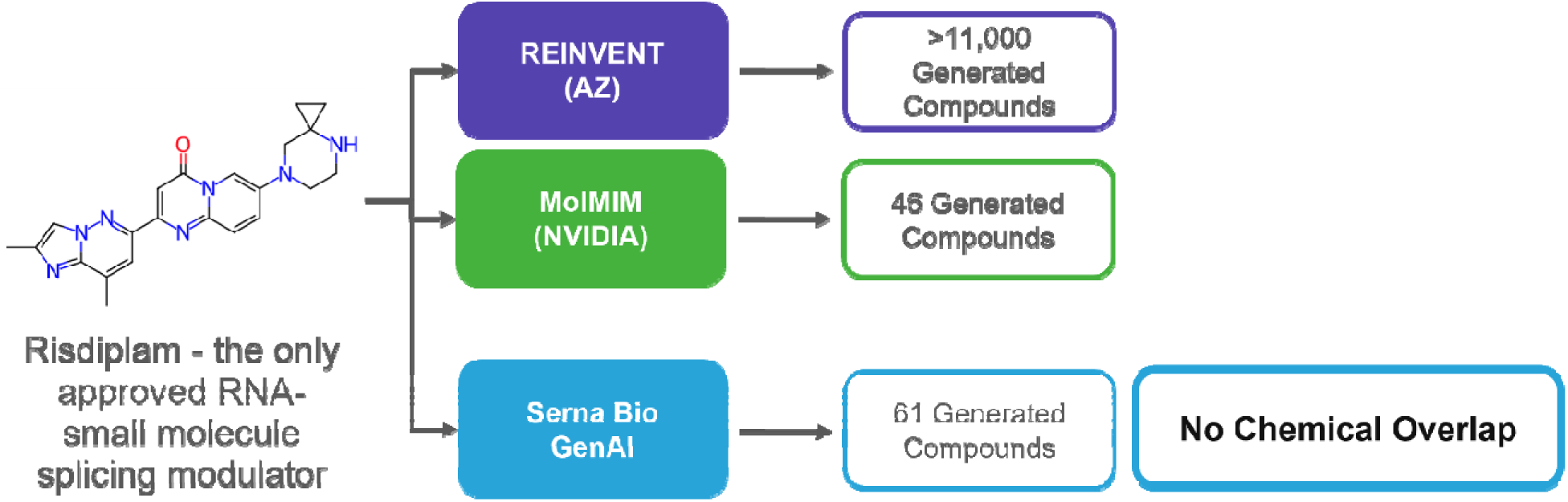
The Serna Bio GenAI designs compounds that are different than those designed by REINVENT and MolMIM when using risdiplam as a seed compound.

**Figure 2.**
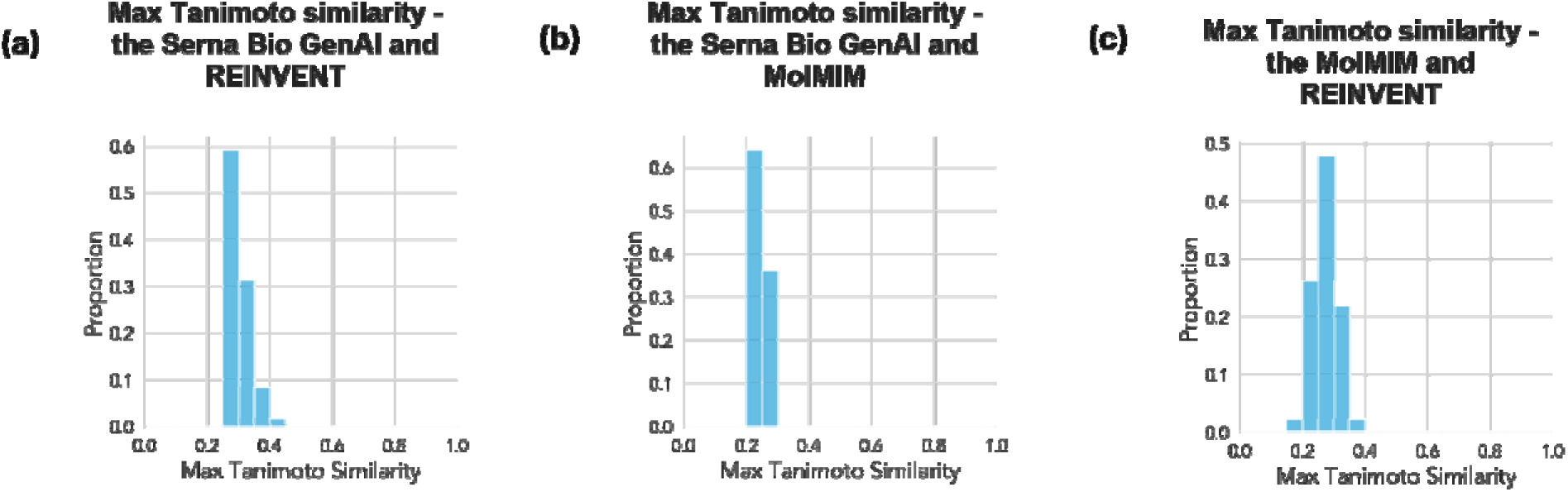
The distribution of pairwise maximum Tanimoto similarity between compounds generated by the chemical generator explored in this work when risdiplam is used as a chemical seed is generally low: (a) the Serna Bio GenAI and REINVENT, (b) the Serna Bio GenAI and MolMIM, and (c) MolMIM and REINVENT.

One goal of the Serna Bio GenAI is to design compounds suitable for hit-to-lead and lead optimization in RNA-targeting drug discovery to try and assist in overcoming SAR cliffs in RNA-targeting drug discovery. Hence, designing compounds with small molecular changes compared to the seed should be considered an advantage, as it allows the drug discovery team to evaluate how these small molecular changes are affecting biological activity. Tanimoto similarity was used to assess the similarity of the risdiplam seed to compounds generated by the different generative chemistry algorithms (Table 1, Fig. 3a). Compounds designed by the Serna Bio GenAI have a mean Tanimoto similarity value to risdiplam of 0.77, with more than 80% of the values being greater than 0.7. None of the REINVENT (mean similarity = 0.15) or MolMIM (mean similarity = 0.18) designs have similarity above 0.7.

**Table 1.**
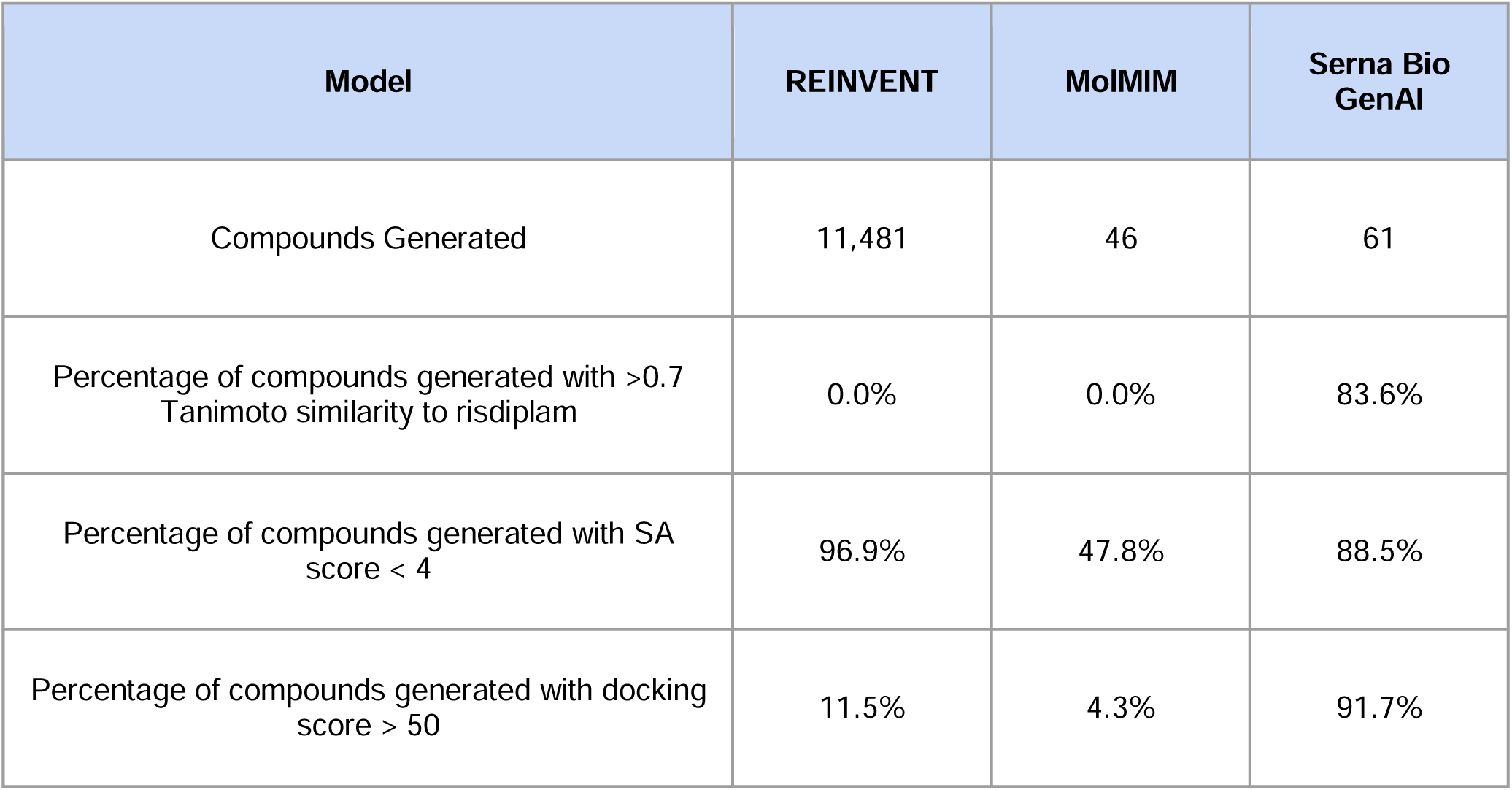
Table comparing the risdiplam analogues designed by REINVENT, MolMIM and the Serna Bio GenAI.

| Model | REINVENT | MolMIM | Serna Bio GenAI |
| --- | --- | --- | --- |
| Compounds Generated | 11,481 | 46 | 61 |
| Percentage of compounds generated with >0.7 Tanimoto similarity to risdiplam | 0.0% | 0.0% | 83.6% |
| Percentage of compounds generated with SA score < 4 | 96.9% | 47.8% | 88.5% |
| Percentage of compounds generated with docking score > 50 | 11.5% | 4.3% | 91.7% |

**Figure 3.**
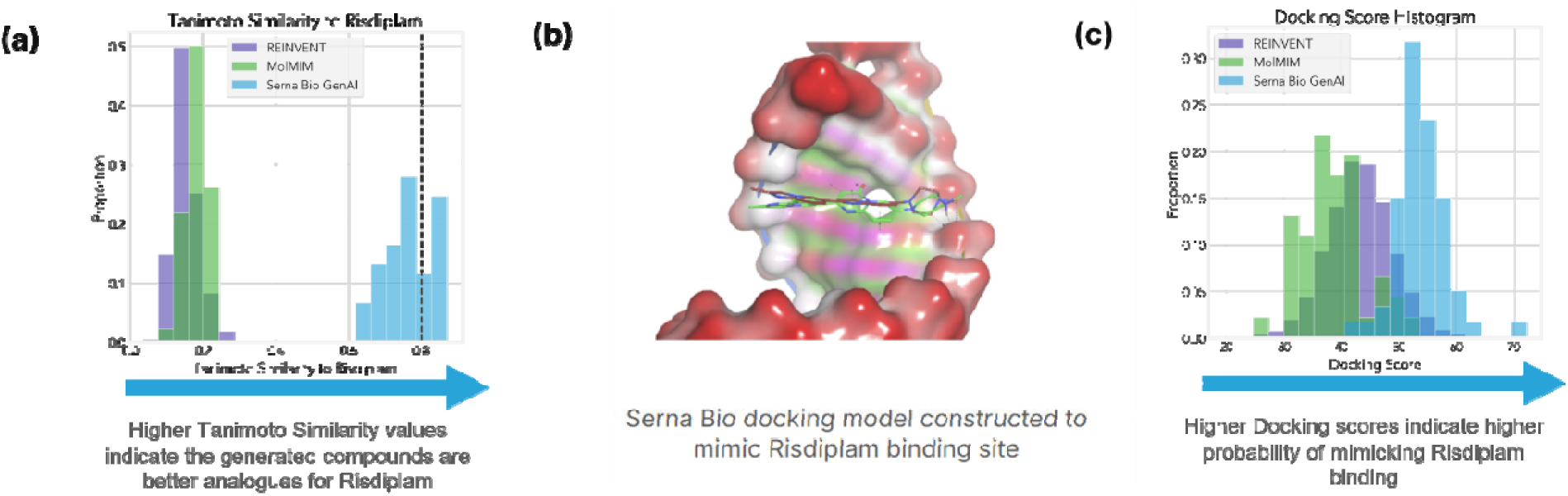
(a) The Tanimoto similarity of compounds generated by REINVENT, MolMIM and the Serna Bio GenAI to risdiplam indicates that only the Serna Bio GenAI is designing similar structural chemistry. A high similarity bar, 0.8, is indicated as a vertical grey dotted line. (b) An image of the SMN2-U1 duplex target used to build our docking study. The experimentally determined position of risdiplam is shown in red and the docked pose by our docking model is shown in green, showing a good match. (c) The docking scores of compounds generated by REINVENT, MolMIM and the Serna Bio GenAI shown in a histogram.

The synthetic accessibility [52] of designed compounds was calculated, and it also highlights a difference between the models (Table 1). SA scores provide simple numerical measures to estimate ease of compound synthesis, and while theoretical synthesizability can be assessed using more modern approaches (including AiZynthFinder [62] and RetroChimera [63]), we believe the simple SA scores are suitable for our comparison here. More than 85% of the Serna Bio GenAI and more than 95% of REINVENT designs have SA scores less than 4, indicating reasonable synthesizability [64], while fewer than 50% of MolMIM designs meet this threshold. Synthesizing the compounds designed by MolMIM may be more challenging.

Finally, a computational docking study using the PDB structure of the SMN2-U1 duplex [53] target of risdiplam was conducted as an *in silico* estimate of compound activity. Risdiplam has been shown to stabilize a specific residue in the 5’ splice site of SMN2 exon 7, strengthening the splice site and resulting in the promotion of exon 7 inclusion [61]. The experimental docking pose of risdiplam in the target is well modelled by our computational model (Fig. 3b) and, when docked, yields a docking score of 48.7. The docking scores of molecules designed by the Serna Bio GenAI, REINVENT and MolMIM support our hypothesis that the fine-tuned generators in our platform are designing a higher proportion of compounds for improving the molecular activity of RNA-targeting small molecules such as risdiplam (Table 1, Fig. 3c). While these results are promising - we acknowledge that *in silico* docking is only a proxy for compound activity, and not equivalent to experimental validation. Furthermore, we acknowledge the chemical similarity of analogues designed by the Serna Bio GenAI to risdiplam likely enables them to fit the SMN2-U1 duplex more effectively than the more diverse chemistry designed by REINVENT or MolMIM.

The combined findings presented above indicate that the Serna Bio GenAI designs synthetically accessible chemistry suitable for the exploration of SAR around risdiplam with promising *in silico* docking scores, making its designs good candidates for our hypothetical RNA-targeting drug discovery campaign.

### The Serna Bio GenAI platform designs different compounds than a human expert chemist and improves compound potency by ∼3X in one DMT cycle

Having demonstrated that the Serna Bio GenAI platform outperforms REINVENT and MolMIM in the *in silico* RNA-targeting exercise using risdiplam as a seed compound, we moved on to compare Serna Bio GenAI-designed compounds to those designed by a human expert medicinal chemist. Validating the AI-designed structures for their synthesizability and in vitro experimental activity represents an important step in understanding the value of such technology to medicinal chemists and drug discovery processes [65]. To do this, we used our platform to design analogues for a Serna Bio RNA-targeting drug discovery campaign. The specific drug discovery campaign in question aims to upregulate the expression of the GLUT1 protein for the treatment of GLUT1 deficiency syndrome (GLUT1-DS). It is important to note that we are designing an entirely novel class of small molecules to enhance protein translation through an interaction with RNA, rather than more classical MOAs such as kinase inhibition.

GLUT1 (SLC2A1) is a membrane protein and is considered an undruggable protein class, a solute carrier [66]. In GLUT1-DS, mutations in one copy of the GLUT1 protein result in insufficient glucose transport to the brain. GLUT1-DS is a severe disorder characterized by intractable seizures, significant movement disorders, and cognitive impairments [67], with no current treatments available for the neurodevelopmental disorders suffered by patients [68]. We aim to discover a small molecule able to bind to the GLUT1 3’UTR. The Serna Bio Target Discovery Platform discovered a novel RNA motif, and by targeting this motif, we can enhance the expression of GLUT1 protein.

The objective of the Serna Bio GLUT1-DS program is to increase the expression of the GLUT1 protein by more than 25% [68]. The change of protein expression for our chemical library of 3,000 compounds was measured in an HCI screen, and 43 primary hits were found, from which we extracted 6 chemical series. The most potent chemical series identified has been shown to bind the 3’UTR RNA Motif in the GLUT1 mRNA identified by the Serna Bio Target Discovery Platform in a biophysical ASMS screening experiment and was advanced to molecular optimization. To evaluate the use of the Serna Bio GenAI platform, we used it to optimize potency for the first DMT cycle. At Serna Bio, we believe in running a head-to-head experimental comparison, and aimed to test ∼20 compounds designed by the Serna Bio GenAI platform and ∼20 compounds designed by a human medicinal chemistry expert.

Using the same hit compounds as seeds, the Serna Bio GenAI designed 411 unique and valid potential compounds for our lead chemical series. 26 compounds were designed independently by a human expert medicinal chemist, and only 2 were found to overlap between these designs and those designed by our GenAI platform (Fig. 4). The maximum and mean pairwise Tanimoto similarities (Fig. 5) between the compounds designed by the Serna Bio GenAI and the human expert medicinal chemist show that there is some chemical similarity between the lists (some pairs have max similarity around or greater than 0.8) and some are more structurally distinct (similarities around 0.2-0.4). These plots are distinctly different, with much higher similarity values, to those comparing the Serna Bio GenAI to REINVENT and MolMIM (Fig. 2). This demonstrates the ability of our generative model ensemble to design material differently from a human expert, albeit exploring a similar section of chemical space around a provided seed, and showcases how chemical design will be directed differently when tools like our platform are available for hit-to-lead and lead optimization. To further consider the novelty of the designed analogues, we searched ChEMBL [23] (Table 2). A lower percentage of Serna Bio GenAI-designed compounds have exact matches, and a similar percentage have highly similar compounds in ChEMBL when compared to Human-designed Compounds, supporting the hypothesis that compounds designed by the GenAI are novel.

**Figure 4.**
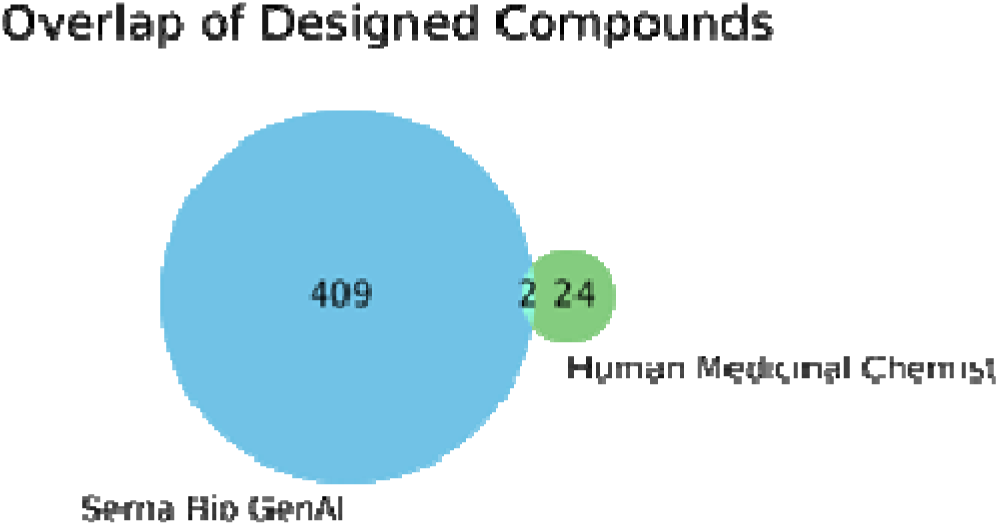
Only 2 compounds were found to be designed by both the Serna Bio GenAI and our human medicinal chemistry expert for the same chemical series using the same initial data.

**Figure 5.**
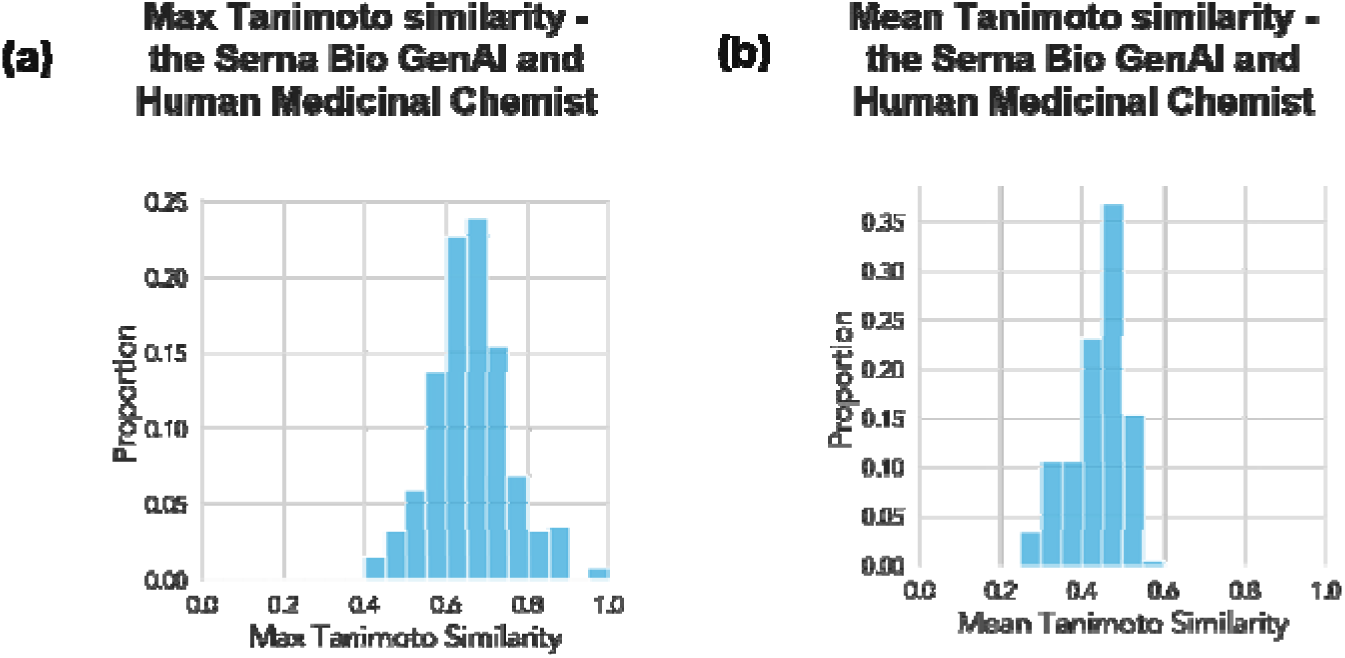
Histograms showing (a) the maximum Tanimoto similarity and (b) the mean Tanimoto similarity between compounds designed by the Serna Bio GenAI and our human medicinal chemistry expert for the same chemical series using the same initial data. These indicate some chemical similarity in the compounds designed by each - particularly when compared to similar comparisons in Figure 2.

**Table 2.** Both Human-designed and Serna Bio GenAI-designed compounds have similar, low percentages of compounds with exact matches or highly similar analogues in ChEMBL.

| Compound Origin | Compounds with exact matches in ChEMBL | Compounds with highly similar analogues (>0.8 Tanimoto similarity) in ChEMBL |
| --- | --- | --- |
| Serna Bio GenAI-designed Compounds | 10/411 (2%) | 95/411 (23%) |
| Human-designed Compounds | 4/26 (15%) | 6/26 (23%) |

As part of the MPO to rank the 411 compounds, we used a set of parameters below: their SA scores [52], druglikeness (using QED scores) [69], and RNA binding probability using a Serna Bio machine learning classifier trained using Serna Bio RNA-small molecule data. SA and QED scores were calculated using RDKit [47]. Serna Bio GenAI-designed compounds were ranked using MPO of these scores (Fig. 6) to guide which are most appropriate for synthesis and testing. Compounds were selected for synthesis and testing by combining the MPO rankings with feedback from our medicinal chemist. When performing chemical synthesis, one human-designed analogue and two GenAI-designed analogues failed to be successfully synthesized, suggesting a slightly lower attrition rate in chemical synthesis for human-designed compounds for this chemical series, but not that the attrition for Serna Bio GenAI-designed compounds is fatal for algorithm deployment in drug discovery.

**Figure 6.**
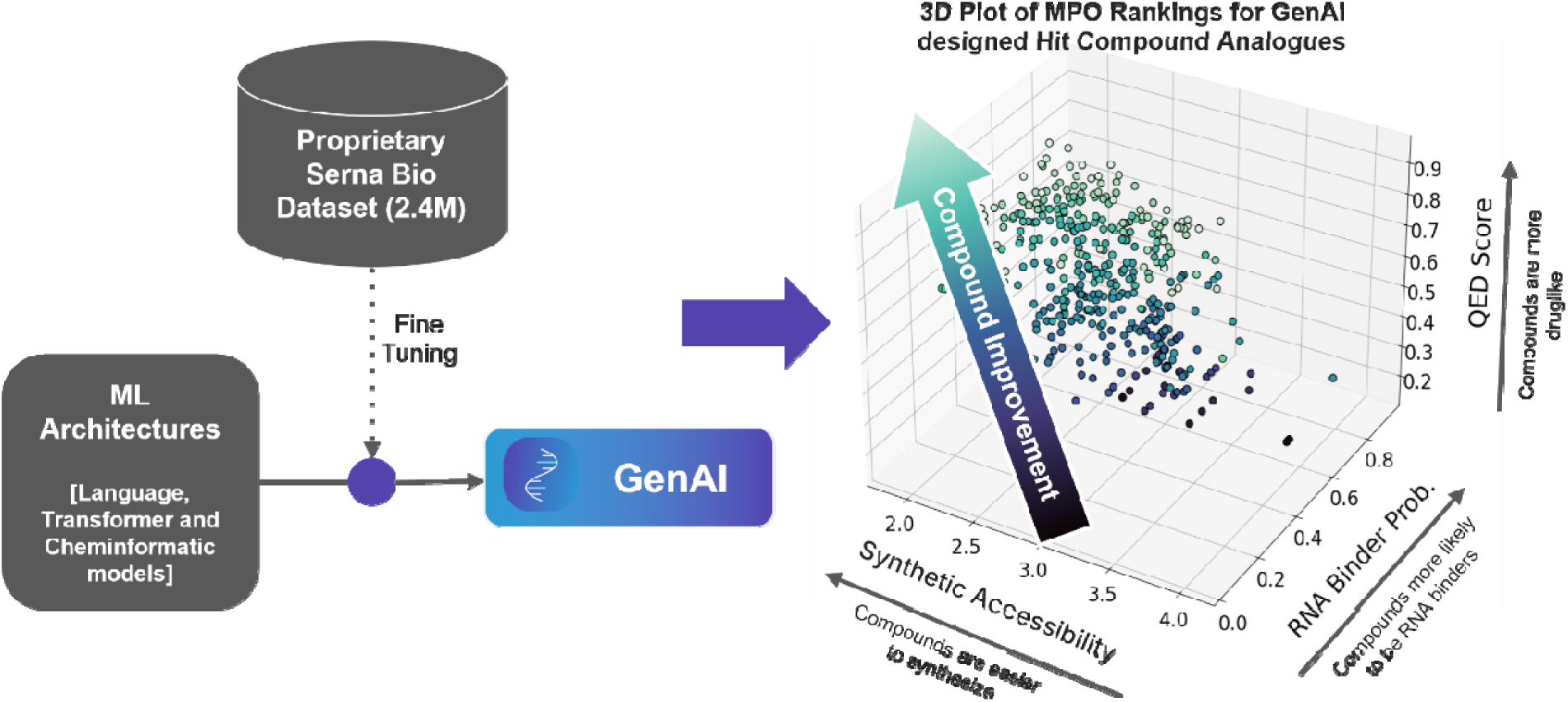
The Serna Bio GenAI consists of generative architectures trained on proprietary Serna Bio RNA-small molecule data coupled with multiparametric optimization to select the best designed chemical analogues. In this example, selections are optimized for higher RNA binding probability using a Serna Bio ML model, high QED scores and low SA scores.

When experimentally tested, the Serna Bio GenAI-designed compounds were found to be the most potent (Table 3), and improved compound potency by ∼3X in a single DMT cycle (Fig. 7, top row).

**Table 3.** A comparison of experimental potency for Human-designed and Serna Bio GenAI-designed compounds from the same chemical series using the same initial data.

| Compound Origin | Compounds Tested | Number with EC25 <10μM | Number with EC25 <2μM | Number with EC25 <1μM |
| --- | --- | --- | --- | --- |
| Serna Bio GenAI-designed Compounds | 22 | 16 (73%) | 9 (41%) | 6 (27%) |
| Human-designed Compounds | 17 | 16 (94%) | 9 (53%) | 2 (12%) |

**Figure 7.**
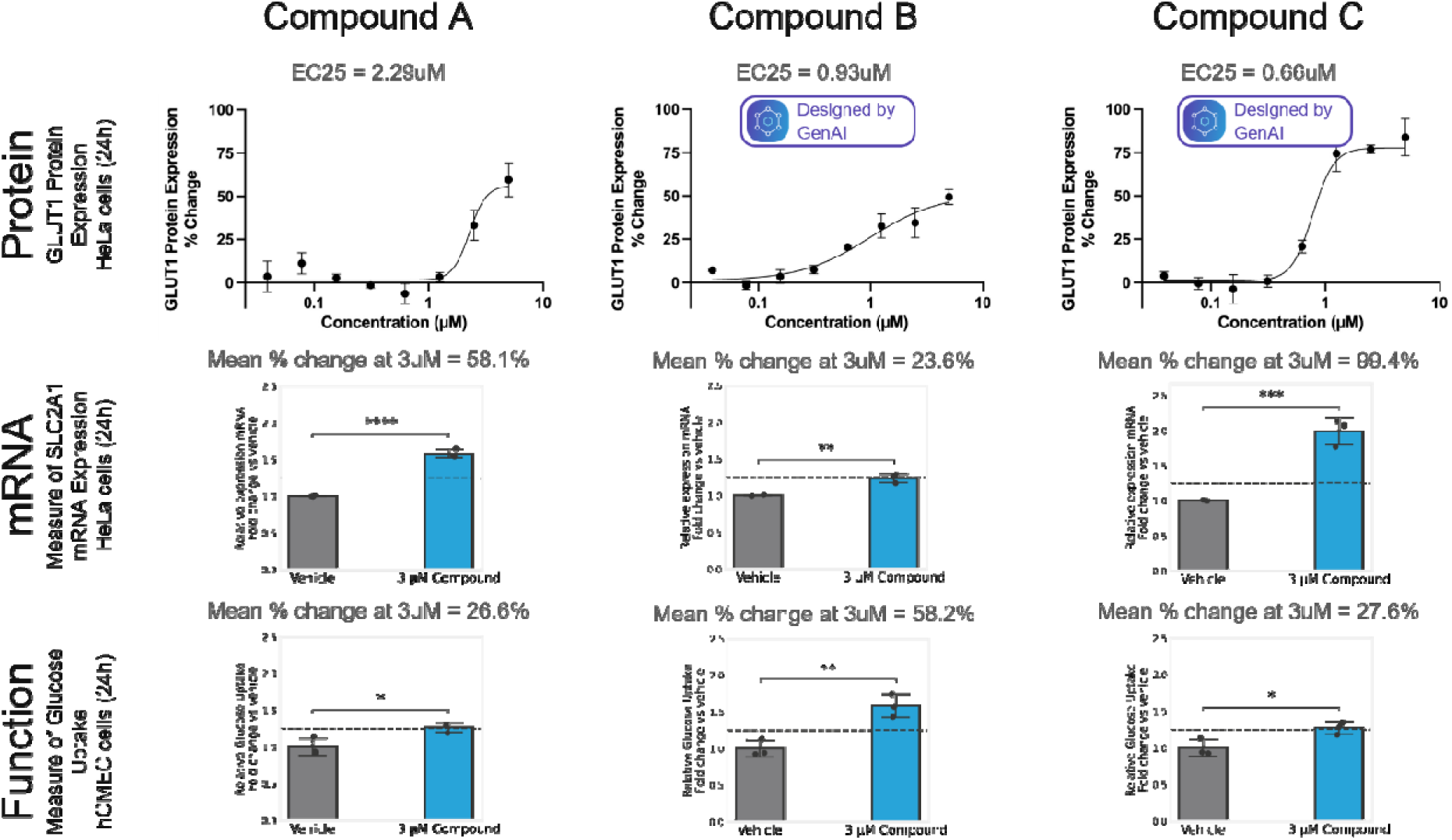
Compounds designed by the Serna Bio GenAI Platform (Compounds B, C) improve compound potency compared to the initial compound series Hit (Compound A). GLUT1 Protein expression concentration-response is presented for each compound with potency being measured by EC25 - the concentration required to result in a 25% upregulation in GLUT1 protein expression. SLC2A1 mRNA expression and glucose uptake tested at 3uM are presented for each compound with mean % change presented as a measure of compound effect. Significance for mRNA expression and glucose uptake change was determined using an unpaired t-test. ns = not significant, *p<=0.05 **p<=0.01, ***p<=0.001, ****p<=0.0001. The dotted line indicates a 25% increase.

### The Serna Bio GenAI platform-designed compounds have confirmed on target activity

To show that Serna Bio GenAI-designed compounds have functional activity beyond the increase in expression of protein, the expression of GLUT1 mRNA and glucose uptake in cells were measured. GenAI-designed compounds previously shown to increase protein expression show a statistically significant increase in GLUT1 mRNA expression at a test concentration of 3µM (Fig. 7, middle row). The GenAI-designed compound C also increases mRNA expression by more than the initial hit compound at 3µM. The two Serna Bio GenAI-designed compounds tested for glucose uptake yielded more than a 25% increase at a test concentration of 3µM (Fig. 7, bottom row).

To demonstrate on-target biophysical target engagement of compounds designed by the Serna Bio GenAI, we conducted ALIS experimental binding studies [44] to our discovered novel RNA-target motif in GLUT1. The initial Hit (Compound A) was a previously identified binder and reconfirmed in a repeat measurement. The Serna Bio GenAI-designed Compound C, with the highest on-target potency of the three compounds featured in this case study, also bound the GLUT1 RNA motif. Compound B was not tested in this experiment. Biophysical binding evidence supports the hypothesis that these compounds are RNA-targeting, act through an interaction with an RNA-motif target, and that the Serna Bio GenAI platform is capable of designing RNA-targeting small molecules.

## Discussion

The presented Serna Bio GenAI platform for RNA-targeting small molecule generation has been compared to state-of-the-art publicly available molecular generators. Our platform was found to be effective at designing chemistry to explore the SAR of a known RNA-targeting seed compound and designing compounds with potential to improve its molecular activity, as demonstrated in an *in silico* docking study - although we acknowledge docking is not equivalent to experimental validation. The Serna Bio GenAI platform’s utility is further demonstrated experimentally in a Serna Bio drug discovery program, with our platform designing chemistry that is different from that designed by a human expert and that competes well in designing hit compounds. Furthermore, the most potent sub micromolar hits were designed largely by our platform, and these hit compounds have confirmed biophysical binding, molecular activity, enhancing the expression of both mRNA and protein levels, and yielding a downstream effect on glucose uptake. The Serna Bio GenAI’s strong performance on RNA-targeting drug discovery tasks, such as our case study GLUT1 program, highlights the importance of building such tools when working on new modalities and tasks with fundamental differences from historical drug discovery. Given RNA-small molecule data, the Serna Bio GenAI platform not only designed different chemistry when compared to other generative algorithms and a human expert medicinal chemist, but that chemistry is useful in advancing an RNA-targeting drug discovery campaign through compound synthesis and experimental testing.

The results of this study demonstrate the value of building specific algorithms for the advancement of RNA-targeting drug discovery programs. However, we do note that our study is limited in scope to the specific results presented for risdiplam and our drug discovery campaign, including the limited sample size of the DMT cycle presented here, and that further evidence is required to truly prove the value and superiority of our GenAI platform. Furthermore, the compared implementations of REINVENT and MolMIM were run using off-the-shelf parameters, and we do not mean to suggest they are not useful chemical generators, or that through model tuning and objective alignment they could not produce more appropriate risdiplam analogues for RNA-targeting drug discovery. Finally, the use of docking scores as a proxy for compound activity is not the same as experimental validation of compound potency.

The results of this study demonstrate the value of generating RNA-small molecule data and using it to fine-tune and train machine learning algorithms for the advancement of RNA-targeting drug discovery programs. As we continue to advance our drug discovery program in lead optimization, we are further improving the Serna Bio GenAI platform to:

- Design and optimize RNA-targeting compounds using targeted local generative and predictive models of our compound series to be more specific to the chemistry and activity desired in the drug discovery program.
- More effectively explore the chemical space around a provided seed compound by incorporating additional generative architectures and fine-tuning these architectures on RNA-targeting small molecule data.
- Optimize other important parameters in drug discovery, including predictive ADME and toxicity modelling, to tackle the multiple facets of lead optimization.

## Supplementary Data

- As a DOC:

○Distributions of physicochemical properties for all compounds in the Serna Bio Dataset, all RNA binder compounds in the Serna Bio Dataset and all compounds in the R-BIND Dataset.
- As an XSLX:

○Tables of Min, Max, Median and Mean of physicochemical properties for all compounds in the Serna Bio Dataset, all RNA binder compounds in the Serna Bio Dataset and all compounds in the R-BIND Dataset.

## Abbreviations

ALIS: Automated ligand identification system
DMT: Design-make-test
GLUT1-DS: GLUT1 deficiency syndrome
HCI: high-content imaging
MLM: Masked language model
MMP: Matched molecular pairs
MOA: Mechanism of action
MPO: Multiparametric optimization
QED: Quantitative estimate of druglikeness
qPCR: Quantitative polymerase chain reaction
SA: Synthetic accessibility
SAMDI-ASMS: Self-assembled monolayer desorption ionization-affinity selection-mass spectrometry
SAR: Structure-activity relationship
TPP: Target product profile

