## Supplementary Data 1 for "Generative Chemistry Platform for Small Molecules Targeting RNA: A Case Study for Chemical Optimization"

*^1^Serna Bio*

*^2^CompChem Solutions Ltd*

*^3^University of Michigan, Biointerfaces Institute*

*

### Supplementary Data

- Figure S1 - Distributions of physicochemical properties for all compounds in the Serna Bio Dataset.
- Figure S2 - Distributions of physicochemical properties for all RNA binder compounds in the Serna Bio Dataset.
- Figure S3 - Distributions of physicochemical properties for all compounds in the R-BIND Dataset.

See also, as an XLSX

- Table S1 - Serna Bio Compounds – Min, Max, Median and Mean of physicochemical properties.
- Table S2 - Serna Bio Binders – Min, Max, Median and Mean of physicochemical properties.
- Table S3 - R-BIND – Min, Max, Median and Mean of physicochemical properties.

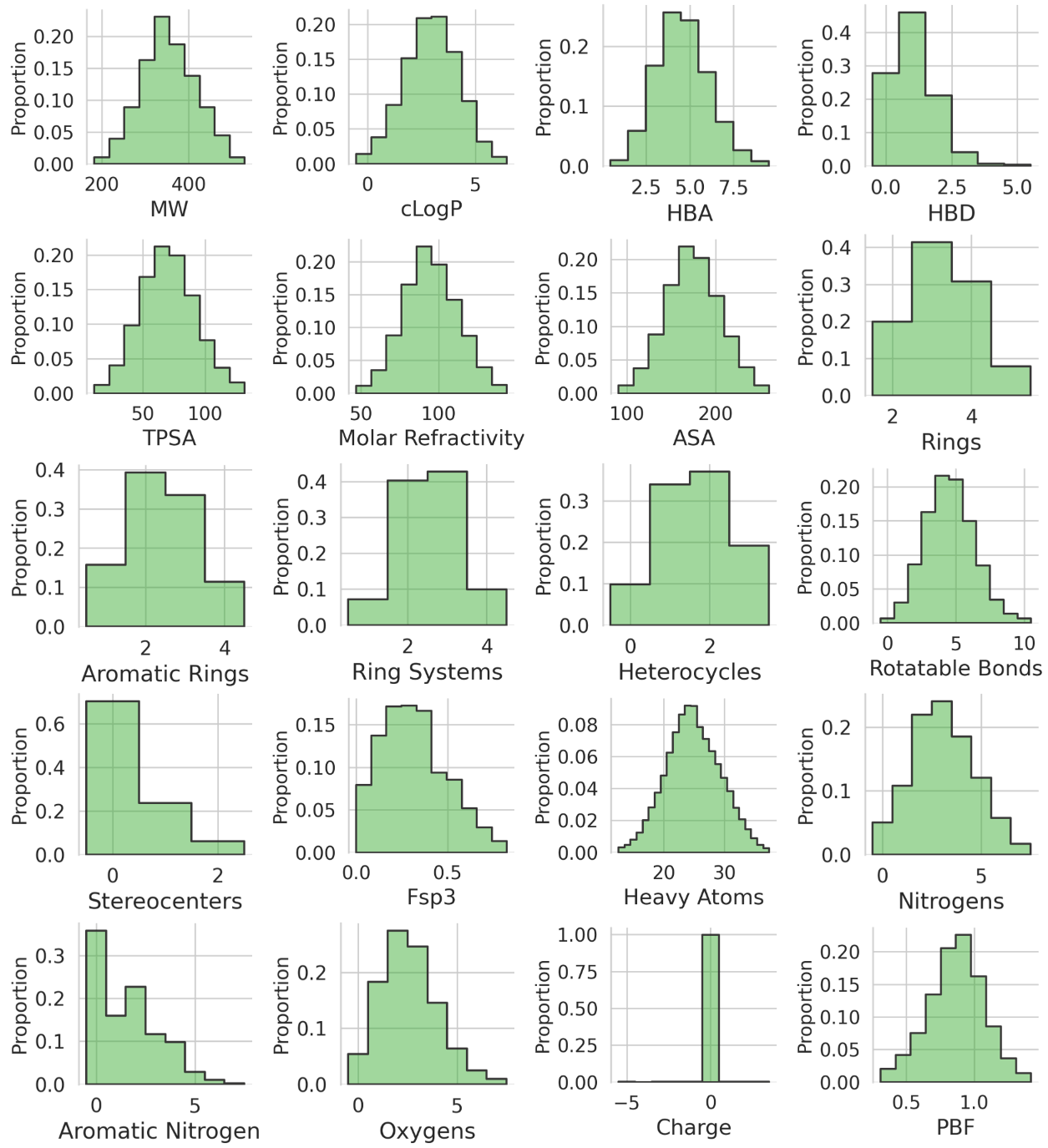

*Figure S1 - Distributions of physicochemical properties for all compounds in the Serna Bio Dataset.*

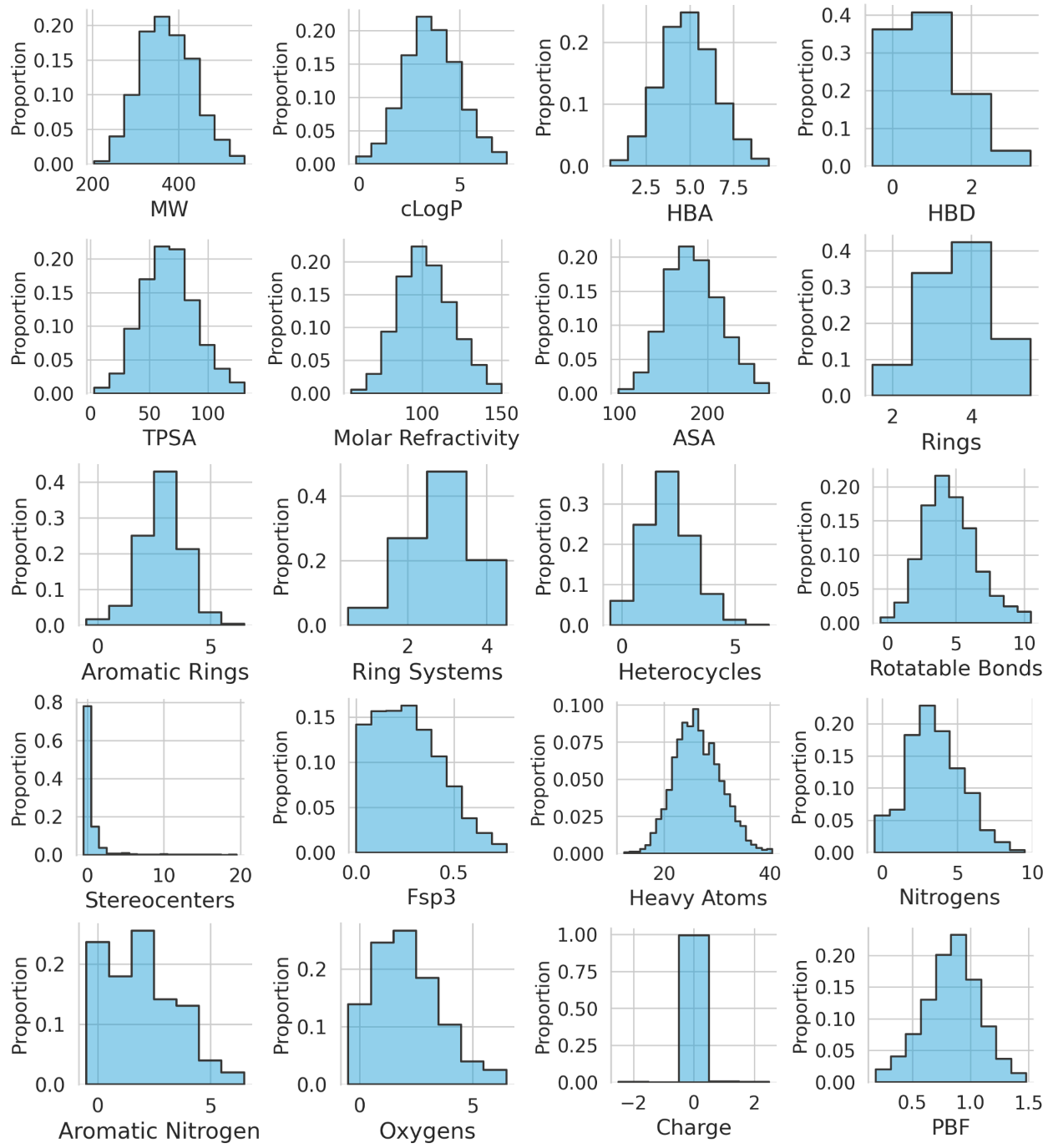

*Figure S2 - Distributions of physicochemical properties for all RNA binder compounds in the Serna Bio Dataset.*

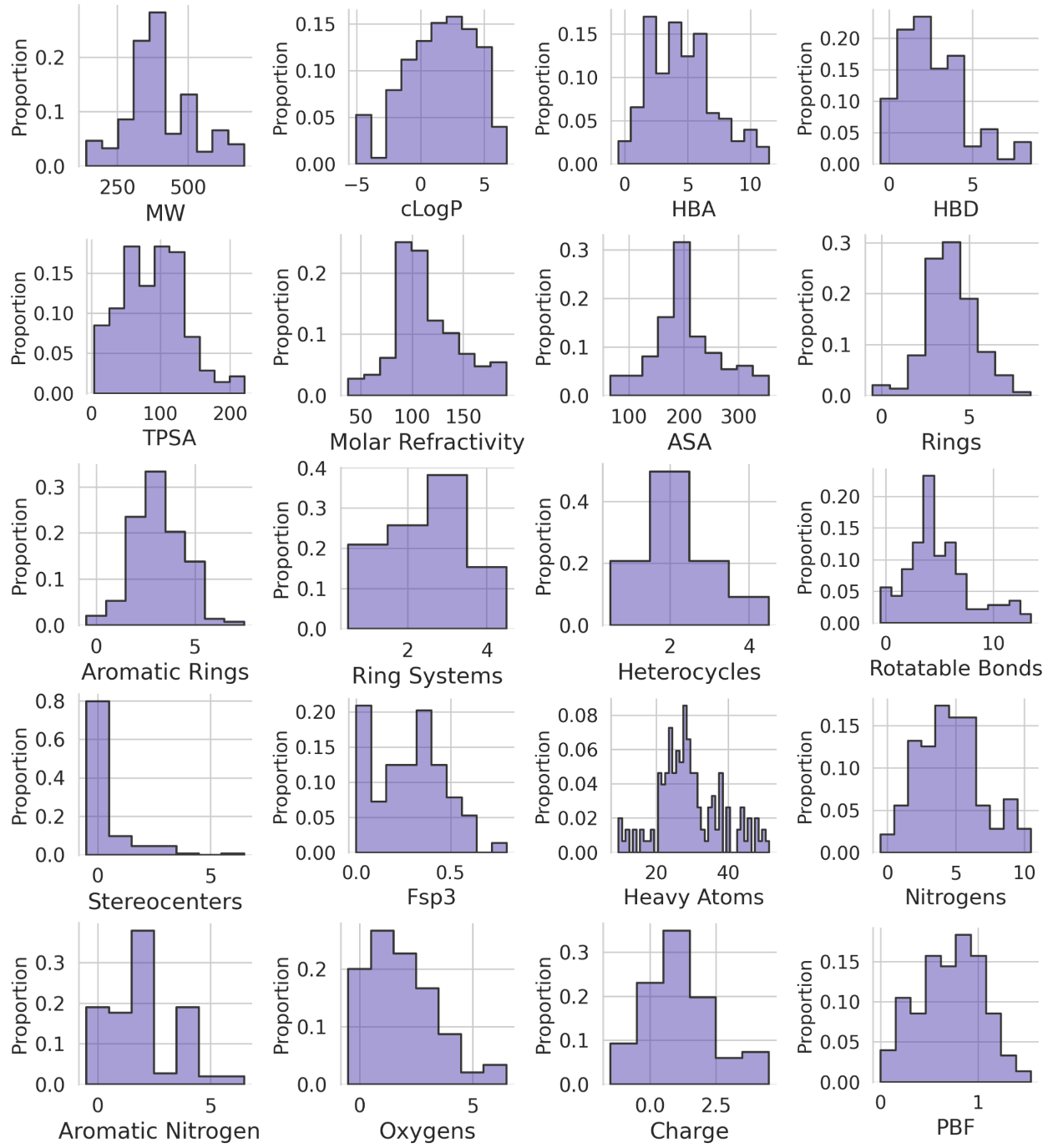

*Figure S3 - Distributions of physicochemical properties for all compounds in the R-BIND Dataset.*
